# In silico analysis of the human milk oligosaccharide glycome reveals key enzymes of their biosynthesis

**DOI:** 10.1101/2022.01.27.477885

**Authors:** Andrew G. McDonald, Julien Mariethoz, Gavin P. Davey, Frédérique Lisacek

## Abstract

Human milk oligosaccharides (HMOs) form the third most abundant component of human milk and are known to convey several benefits to the neonate, including protection from viral and bacterial pathogens, training of the immune system, and influencing the gut microbiome. As HMO production during lactation is driven by enzymes that are common to other glycosylation processes, we adapted a model of mucin-type GalNAc-linked glycosylation enzymes to act on free lactose. We identified a subset of 11 enzyme activities that can account for 206 of 226 distinct HMOs isolated from human milk, and constructed a biosynthetic reaction network that identifies 5 new core HMO structures. A comparison of monosaccharide compositions demonstrated that the model was able to discriminate between two possible groups of intermediates between major subnetworks, and to assign possible structures to several previously uncharacterised HMOs. The effect of enzyme knockouts is presented, identifying β-1,4-galactosyltransferase and β-1,3-*N*-acetylglucosaminyltransferase as key enzyme activities involved in the generation of the observed HMO glycosylation patterns. The model also provides a synthesis chassis for the most common HMOs found in lactating mothers.

## Introduction

Human milk, aside from its value as an optimal source of food for the new-born infant, has increasingly been shown to have many additional benefits in promoting development and providing protection from disease. Human milk oligosaccharides (HMOs) are the third most abundant constituent of milk, after lactose, and lipids, with total mean concentrations ranging from 4–30 g/L [1]. The majority of these complex oligosaccharides are typically based on free lactose. Although indigestible by the neonate, HMOs provide several benefits to it, including antimicrobial action [2, 3], protection from viral pathogens [4–7], promotion of a healthy gut microbiota [8, 9] and the development of the immune and nervous systems [10– 12]. HMOs may also play a role in preventing allergic reactions during childhood [13].

A feature unique to human milk is the heterogeneity of its HMO population, when compared to milks from other species, such as bovine, which are lower in both quantity and structural diversity [14]. Anti-adhesive properties of HMOs rely on competitive inhibition with pathogens for host receptors, as is the case, for example, with the finding that mono- and difucosylated oligosaccharides inhibit the binding of cholera toxin to glycosylated receptors of human epithelial cells [15]. By presenting a wide variety of epitopes, the human milk oligosaccharide is able to mask the newborn from such toxins, while simultaneously promoting a colonisation of beneficial gut flora of Bifidobacterial species [16].

In consequence, there has been much interest towards the synthesis of HMOs industrially, for use as probiotics, for which a number of approaches have been used, including direct chemical syntheses, use of recombinant glycosyltransferase (GT) activities [17]; transglucosidase reactions, in which glycohydrolases act in reverse to attach monosaccharides to oligosaccharides [18–20]; and through reconstruction of Leloir-type [21] pathways to synthesise the preferred HMO substrate of bifidobacteria, lacto-*N*-biose I [22]. Although little is known as to the actual enzyme activities expressed in the lactating mammary epithelium [23], some insights can be gleaned from analysis of the more than 200 HMO structures already characterised, many of which have been employed as model substrates of glycosidases and glycosyltransferases (for example, [24]).

Based on the observation that mammary gland is secretory in nature, with mucosal-like properties [25], a view that has also been supported by the murine case [26], we assumed that some of the enzymes involved in mucin biosynthesis would also be active in the production of HMOs. Our purpose in this article is to apply an adapted model of the enzymes involved in mucin-type (GalNAc-linked) O-glycosylation to act on free lactose, and to validate it against a large population of experimentally characterised HMOs, and subsequently predict a minimal biosynthetic reaction network. In doing so, we extend the previous work of GlycomeSeq [27], an existing *in silico* model of HMO biosynthesis developed as an aid in sequencing milk glycans and that was missing sialylated structures. We also complement the metabolic reconstruction approach of Bao *et al*. (2020) [28] that modelled and scrutinised the activity of ten glycosyltransferases in the synthesis of HMOs detected in cohorts including both secretor and non-secretor healthy mothers. The present article describes the model and the biosynthesis simulations performed on HMOs.

## Results

### Model description

We developed a model of the enzymes of O-linked glycosylation [29] to allow their action on free lactose and predict possible human milk oligosaccharide products. The model uses a formal-grammar based approach to apply regular-expression based rules to model the transfer of monosaccharides, represented by a single-letter code, from activated sugars (sugar-nucleotide donors) to an oligosaccharide acceptor. The transformation rules are classified into a number of discrete types: extension, decoration, branching and termination (see Methods).

We compiled a library of 226 lactose-based human milk oligosaccharides from a variety of sources [23, 30–38], as shown in Supplementary Table S1. The HMOs were found to be composed of the five monosaccharides, L-fucose, D-galactose, D-glucose, *N*-acetyl-D-glucosamine, with a small number with 6-*O*-sulfated residues (4) and a single occurrence of *N*-acetyl-D-galactosamine. A single-letter code was used to denote these monosaccharide units, as shown in Table 1. Free lactose, β-D-Gal*p*-(1,4)-D-Glc*p*-ol, is represented by [L4G]. Branched and decorated oligosaccharides are similarly represented by using bracketed notation, for example the monofucosylated HMOs 2′-FL and 3-FL are represented as [[f2]L4G] and [L4[f3]G].

**Table 1.**
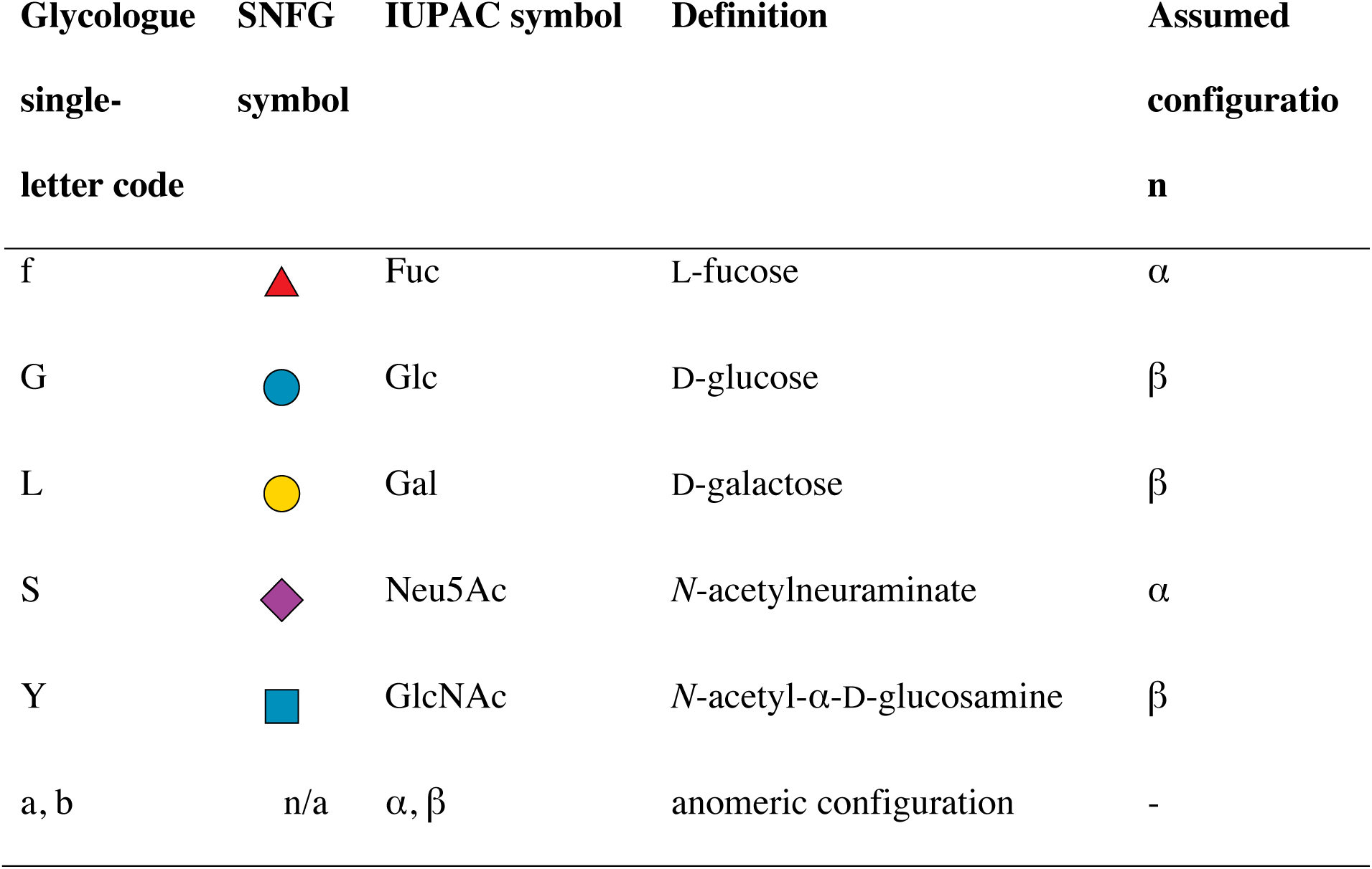
Monosaccharide symbols. Definition of monosaccharide units of HMOs represented as single-letter codes in Glycologue, symbols in the Symbol Nomenclature For Glycans (SNFG) notation [39], IUPAC symbols. For example, *N*-acetylgalactosamine (GalNAc) is represented by Glycologue as ‘V’, and sulfate by the lowercase ‘s’.

| Glycologue<br>single-letter code | SNFG<br>symbol | IUPAC symbol | Definition | Assumed<br>configuration<br>n |
| --- | --- | --- | --- | --- |
| f | | Fuc | L-fucose | $\alpha$ |
| G | | Glc | D-glucose | $\beta$ |
| L | | Gal | D-galactose | $\beta$ |
| S | | Neu5Ac | *N*-acetylneuraminate | $\alpha$ |
| Y | | GlcNAc | *N*-acetyl-α-D-glucosamine | $\beta$ |
| a, b | n/a | $\alpha, \beta$ | anomeric configuration | - |

A table of the most commonly occurring HMOs, ranked according to their frequency of occurrence in the cited literature, is available online at https://glycologue.org/m/sample.php.

### Enzymes and reactions

The enzymes of the model are shown in Table 2, numbered **1**–**11**, ordered by EC number, comprising two galactosyltransferases (enzymes **1** and **6**), three are *N*-acetylglucosaminyl- transferases (**4, 10, 11**), with three fucosyltransferases (**2, 3, 5**) and three siayltransferases (**7**– **9**).

**Table 2.**
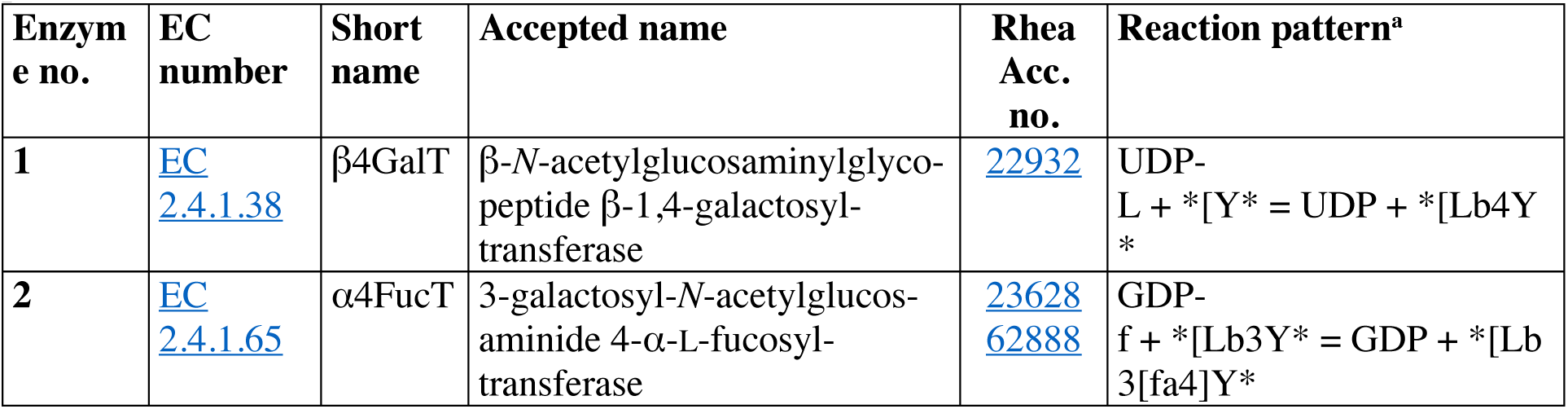
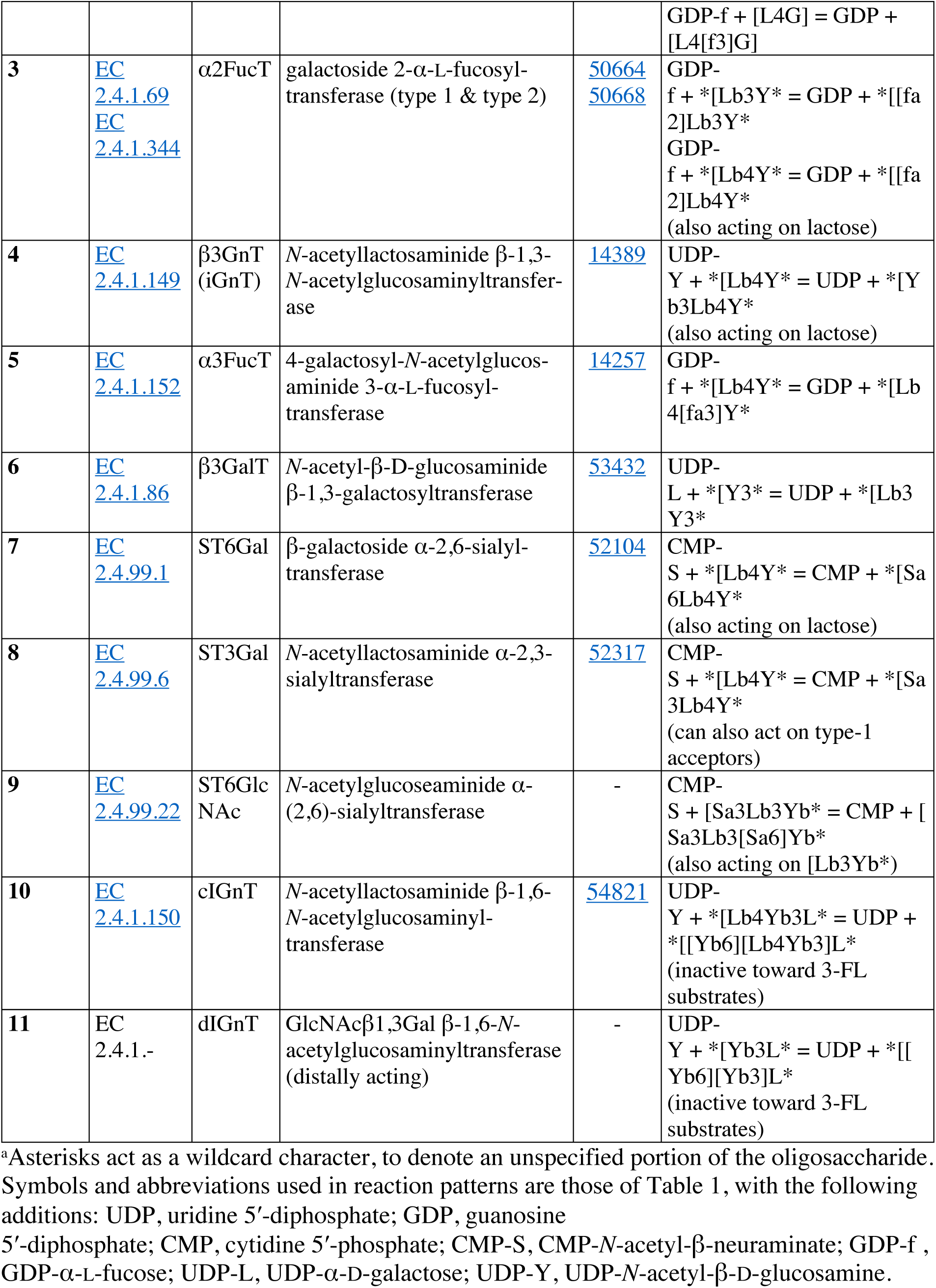
Enzymes of the model. Proposed enzymes of HMO biosynthesis and the corresponding reaction definition in Rhea (accession number) as well as Glycologue reaction patterns.

As shown in Fig. 1 (Linkage types), HMOs, in common with mucin-type O-glycans and other glycoconjugates, possess non-reducing termini that are based on Gal-β1,3-GlcNAc and Gal- β1,4-GlcNAc motifs, denoted type 1 and type 2 [40, 41], respectively, which are formed by the actions of enzymes **6** and **1** on a structure terminating in GlcNAc.

**Figure 1.**
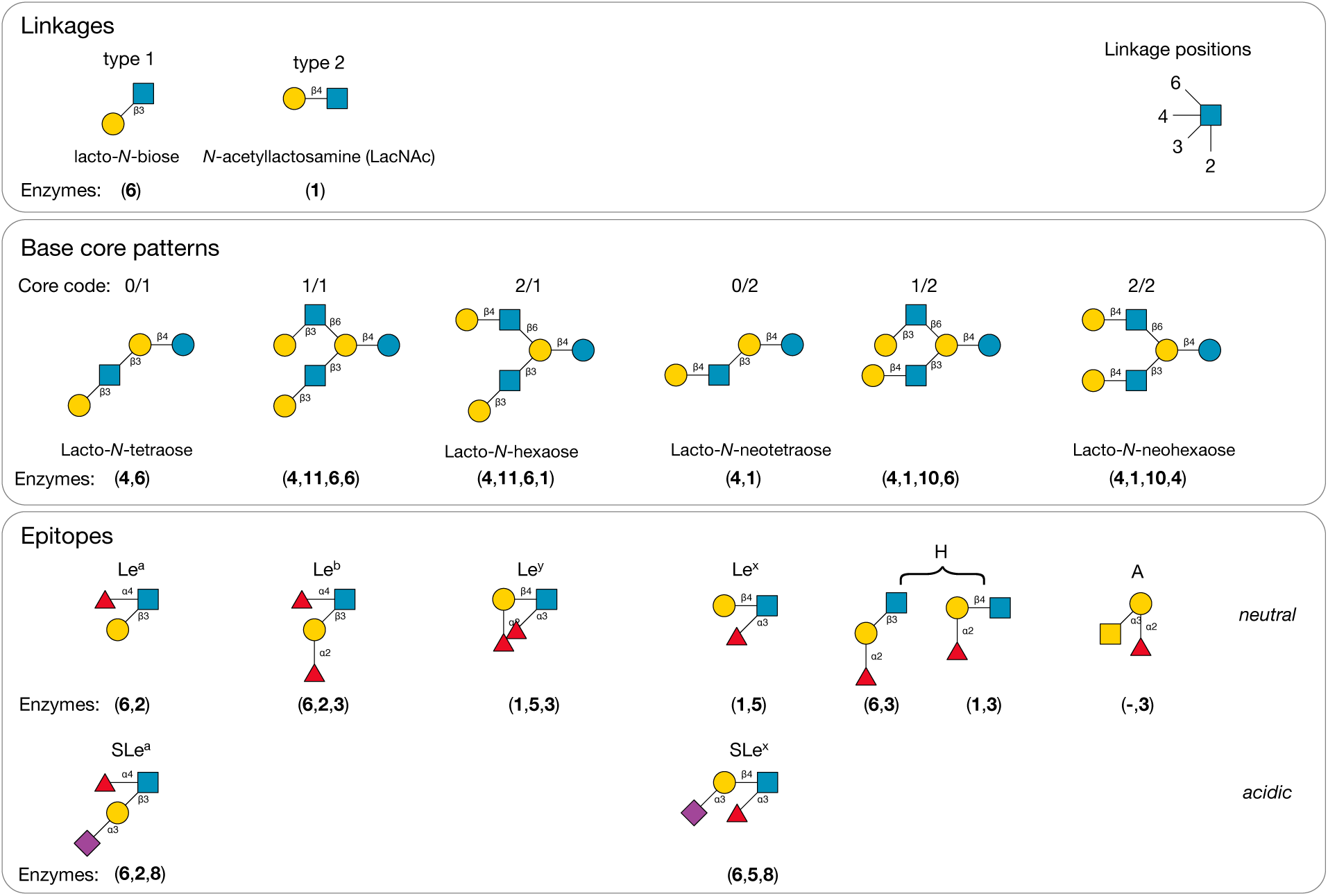
Structural motifs common to HMOs. Core structures are defined as lactose-based oligosaccharides that contain only hexose sugars, Glc and Gal, and the *N*- acetylhexosamine sugar, GlcNAc. Not all cores and epitopes are found within the experimentally determined HMO structures considered by this study. Base core patterns are defined as *a*/*b*, where *a* and *b* refer to the type-1 or -2 linkage of the terminal Gal, either to the 6-linked (*a*) or the 3-linked GlcNAc (*b*) of an I-antigen branch appearing on lactose. Proposed sequences of enzymes (indices of the enzymes in Table 2) involved in the biosynthesis of each motif are displayed beneath its structure. A complete set of core structures is in Supplementary Table S2.

These basic determinants are named lacto-*N*-biose (LNB) and *N*-acetyllactosamine (LacNAc). De-galactosylated HMOs, terminating in GlcNAc, are rare, with only two representatives in Supplementary Table S1: GlcNAcβ1-3Galβ1-4Glc (LNTri II) [42], and one synthesised *in vitro* by Prudden et al. [31], GlcNAcβ1-3Galβ1-4GlcNAcβ1-3Galβ1- s4GlcNAcβ1-6 (Neu5Acα2-6Galβ1-4GlcNAcβ1-3)Galβ1-4Glc.

The smallest model network employing all of the activities of the enzymes in Table 2, is shown in Fig. 2.

**Figure 2.**
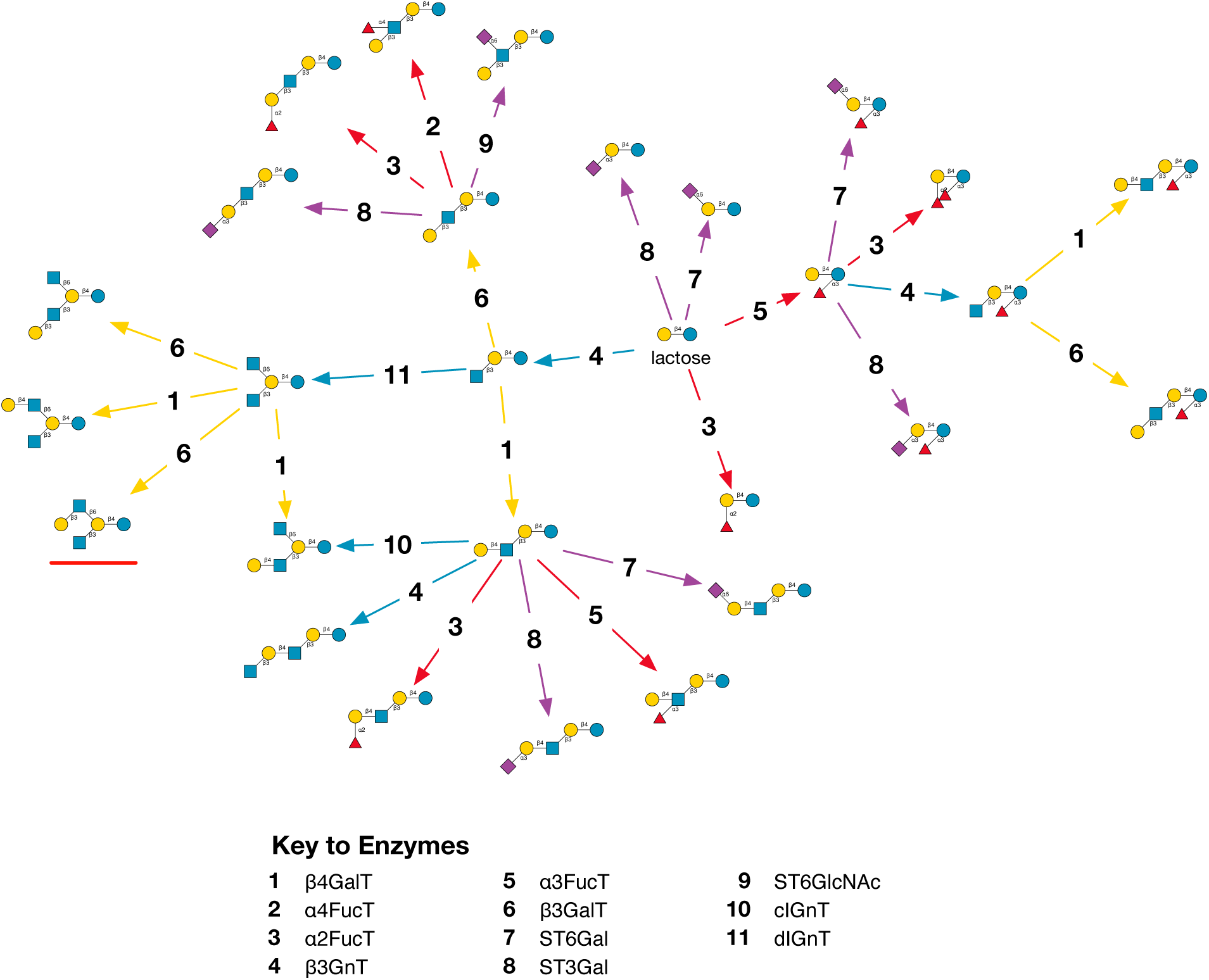
Reactions of the model. A reaction network generated by three iterations of the enzyme simulator, starting from lactose. Enzymes reactions are represented arrows leading from an acceptor substrate to an acceptor product, and coloured according to the type of monosaccharide transferred: GlcNAc (blue), Gal (yellow), Fuc (red), Neu5Ac (purple). For simplicity, donor substrates and products are omitted. The structure underlined in red was not found among the observational data set (Supplementary Table S1).

### Bifunctional enzymes

Glycosyltransferases (GTs) are multi-specific, acting on a range of acceptors according to a recognition motif. It is known that many enzymes, including GTs, can display secondary activities, a behaviour sometimes called enzyme promiscuity [43]. Several such are included in the model. For example, in Table 2 possess the α2FucT (**3**) combines the activities of the type 1 and type 2 galactoside 2-α-L-fucosyltransferases, which transfer fucose to either lactose, to form 2′-FL, or lacto-*N*-biose or LacNAc termini. The activities of the other fucosyltransferases, α3FucT and α4FucT, are recognised separately; however, since α4FucT bears a 3-α-fucosyltransferase towards the glucose of free lactose [44], this was included as a secondary activity of enzyme **2**.

### Atypical sialylation

The ST6GlcNAc gene family is not found in humans [45], nevertheless several studies have demonstrated Neu5Ac 6-linked to the GlcNAc of lacto-*N*-biose, as for example in LST b and DS-LNT [32–34, 38]. Prudden and co-workers were able to synthesise the latter by means of the ST6GALNAC5 [31], thus providing evidence of a potential candidate for the primary activity of EC 2.4.99.22 (**9**) in humans. Although the enzyme from rat liver is able to act on asialylated termini ([L3Y) [46], these are not substrates for ST6GALNAC5 [47]. A suitable candidate for the secondary activity of **9** remains to be determined, thus its existence is inferred.

### Branched HMOs

The model incorporates two separate I-branching (beta-1,6-N-acetylglucosaminyltransferase) enzyme activities, one centrally acting and the other distally acting, relative to the reducing end of the oligosaccharide. The central activity, named cIGnT, is that of EC 2.4.1.150, which catalyses the reaction

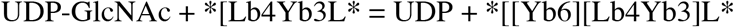

having a preference for oligosaccharides with type 2 termination [48], i.e., Gal β-1,4-linked to GlcNAc. The second IGnT enzyme, dIGnT acts on the predistal galactose before a terminal GlcNAc, and is a secondary activity of the mucin-type core-4 forming enzyme, C2/4GnT [49, 50]. Based on the observation that none of the characterised HMOs that were 3-fucosylated on the base glucose were branched, an additional assumption of the model was that the I-branching enzymes **10** and **11** are inactive towards these substrates.

### Performance of the simulator

Starting from lactose, with all 11 enzymes active, 206 of the 226 HMOs in Supplementary Table S1 were predicted *in silico*, a prediction rate, or “coverage”, of 91.1%. When considering the HMOs common to more than one study, 85 out of the 89 structures listed in the online table were obtainable in simulations, for a 96% coverage. Owing to a combinatorial explosion in the number of structures formed [51, 29], simulations were limited to a user-defined maximum number of GlcNAc residues incorporated into HMO acceptor-products. The number of structures appearing with each iteration of the enzyme simulator increased logistically with iteration number (Fig. 3), when HMOs were limited to between 3 and 6 GlcNAc.

**Figure 3.**
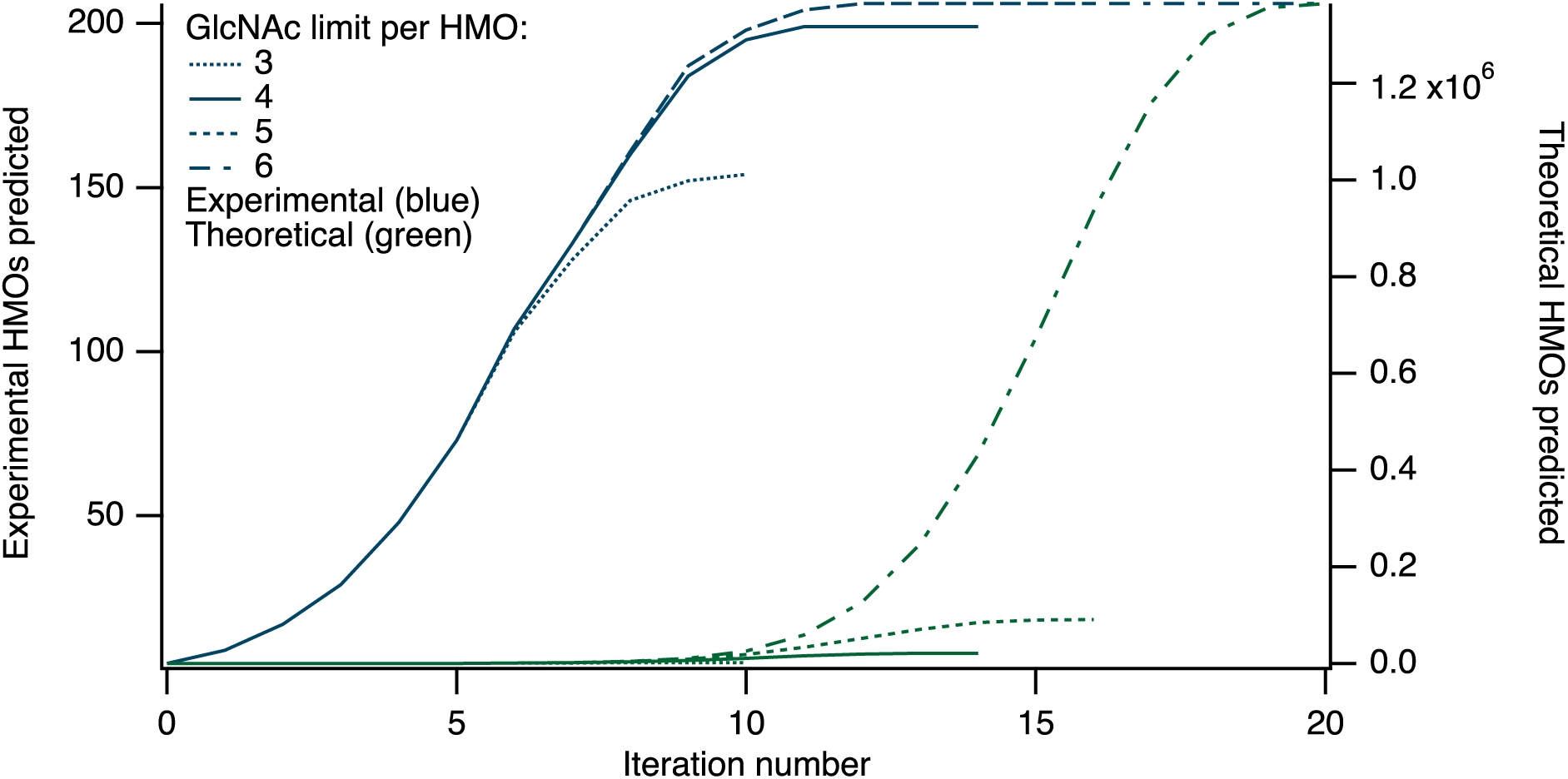
Growth of HMOs predicted by the simulator. Numbers of experimental (left axis) and theoretical (right axis) HMOs predicted, with increasing iteration number, for simulations limited to between 3 and 6 GlcNAc residues per oligosaccharide. At the final point on each curve, reaction networks were closed, with no further structures being added at higher iteration values.

Reaction networks that eventually closed, with no further products being added at higher iteration numbers (see Methods). The maximum number of observed HMOs found in the simulations, while minimising the total number of theoretical structures, was found to occur at iteration 11, when the maximum number of GlcNAc residues per HMO was limited to four.

### Networks and HMO classification

At 11 iterations of the method, and with a limit of 4 GlcNAc residues incorporated per acceptor, the simulator generated 10,821 unique HMO structures, shown in the reaction network in Fig. 4A, where each node is coloured according to the base core structure of the HMO it represents, as shown in Fig. 1 (Base core patterns).

**Figure 4.**
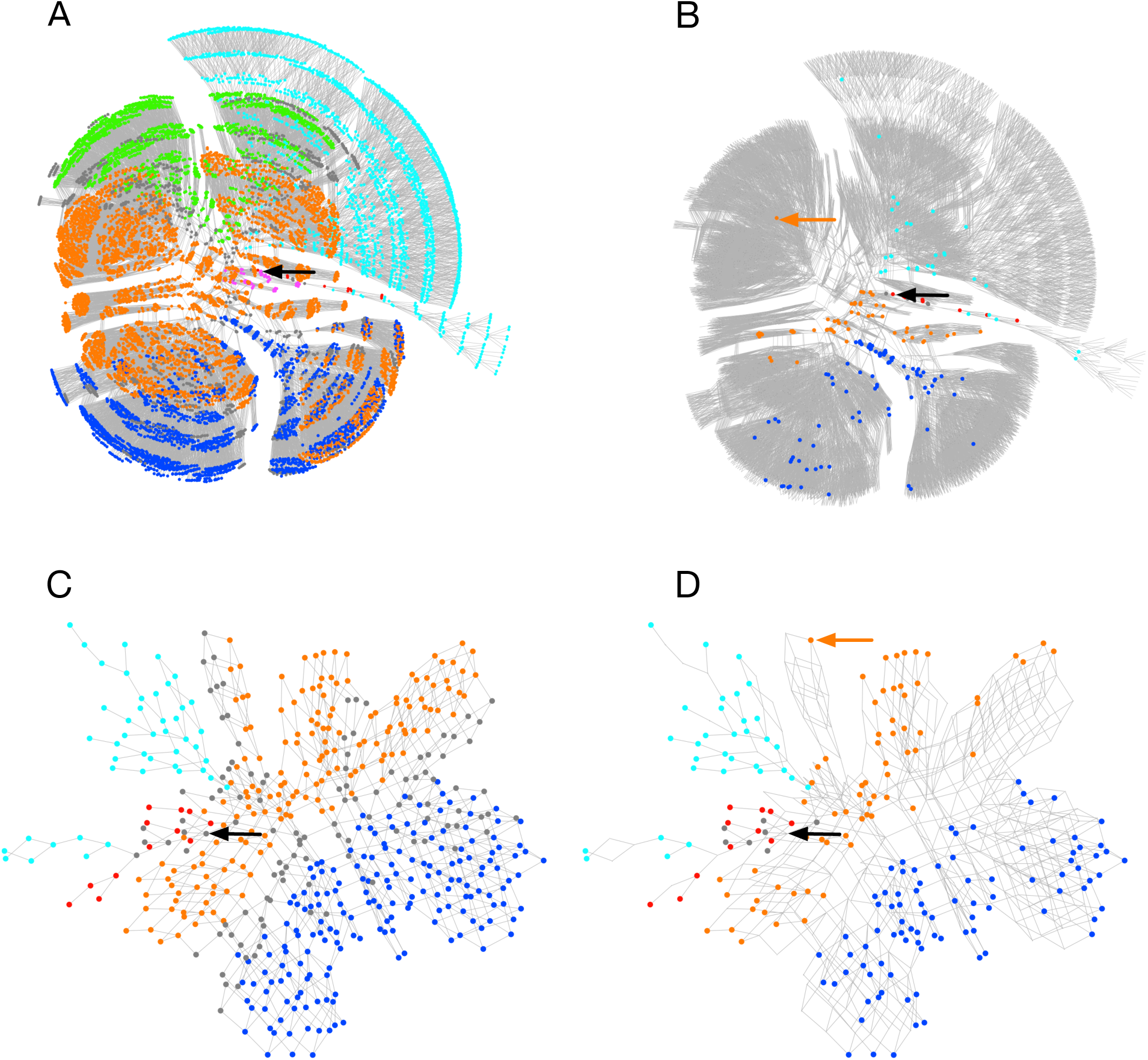
Calculated biosynthetic pathways of human milk oligosaccharides. A. Simulated network of the 11 enzymes of Table 2, limiting to 11 iterations, with each HMO limited to a maximum of 4 GlcNAc residues. **B**. As A, with 195 experimentally characterised HMOs highlighted and coloured, with theoretical structures shown in grey. **C**. A minimal reaction network leading to the population of 206 HMOs from the library of observed HMOs (Supplementary Table S1). **D**. As C, with only the experimentally observed HMOs highlighted. The position of the starting substrate, lactose, is indicated by a black arrow. Nodes are coloured according to the base core configuration given in Fig. 1: magenta (1/1), blue (2/1), green (1/2), orange (2/2), red (0/1), cyan (0/2), grey (other, unclassified). The location off *inverse*-LNnD with networks B and D is indicated by an orange arrow. Networks were drawn in Tulip [52] using a stress-minimization layout algorithm.

Chen [30] identified 16 distinct core HMO structures, non-fucosylated and neutrally charged, composed only of alternating galactose and *N*-acetylglucosamine units and the glucose moiety of lactose. Although lactose is not itself classed as an HMO, under this classification it is Core I. Urashima *et al*. [23] extended this number of core structures to 19 (I–XIX). Among the HMOs considered in the present study, we proposed an additional five novel cores to those of the latter classification. The full set of HMO core structures found is given in Supplementary Table S2.

Since both earlier classifications provided a large number of distinct sub-populations that were found not to correlate spatially in layouts of the biosynthetic networks (Fig. 4), it made their interpretation difficult. For easier visualisation of the networks, we adopted a simpler classification system based on the initial actions of β3GnT (**4**), followed by those of the two galactosyltransferases, β4GalT (**1**) and β3GalT (**6**), and then the branching *N*- acetylglucosaminyltransferases, cIGnT (**10**) and dIGnT (**11**). This approach divides the oligosaccharides into six classes: two that are linear (types 1 and 2, terminating in β-1,3-Gal and β-1,4-Gal, respectively), and four that combined the actions of cIGnT and dIGnT with subsequent extension by β3GalT and β4GalT. We denoted these four branching possibilities by *a*/*b*, where *a* signifies the type of the 6-linked GlcNAc (upper branch), and *b* the 3-linked GlcNAc (lower branch), leading to four combinations of types 1 and 2, namely, 1/1, 1/2, 2/1 and 2/2. In common biochemical nomenclature, 0/1 is lacto-*N*-tetraose (LNT; Core II), 2/1 is lacto-*N*-hexaose (LNH; Core V), 0/2 is lacto-*N*-neotetraose (LNnT; Core III) and 2/2 is lacto- *N*-neohexaose (LNnH; Core VI).

Fig. 4B highlights the observed structures, and their distribution through the network. All regions of the main network have observed counterparts, with the exceptions of the 1/1 (magenta) or 1/2 (green) branching pattern. For the 206 observed HMOs predicted by the model, a minimal biosynthetic network was constructed by reversing the enzymes of glycosylation and compiling a network of all of the reversed-reversed reactions leading from lactose to observed products. The resulting reduced network is shown in Fig. 4C and D, coloured according to the base-core scheme of Fig. 1. The network with 514 HMOs (nodes), including intermediates, and 966 distinct reactions (edges). In Fig. 4D, only the nodes matching the predicted, experimentally observed, HMOs of Supplementary Table S1 are coloured (206 nodes). On account of the multiantennary nature of many HMOs (167 of the 226 HMOs studied bear at least one β-6-GlcNAc residue), multiple routes to the same product are possible, which results in a lattice-like appearance of the networks. This is also seen in other studies of glycosylation reaction networks, including those of HMOs, N-linked and O-linked glycans [28, 29, 53].

A review of HMO concentrations in mature human milk, pooled from 57 studies published between 1966 and 2020, identified the 15 oligosaccharides that were in greatest abundance [54]. All 15 HMOs were predicted within the model, and a biosynthetic network was constructed (Supplementary Figure S1), in which all of the enzymes except the cIGnT activity (**10**) were used. It was observed that none of the most abundant HMOs were of the base-core 2/2, but were linear 0/1, 0/2, or branched 2/1, according to the classification scheme employed here.

### Novel cores/Delayed branching

A possible biosynthetic pathway of eighteen of the core structures I–XIX, along with the structures of the five novel cores, is shown in Fig. 5A.

**Figure 5.**
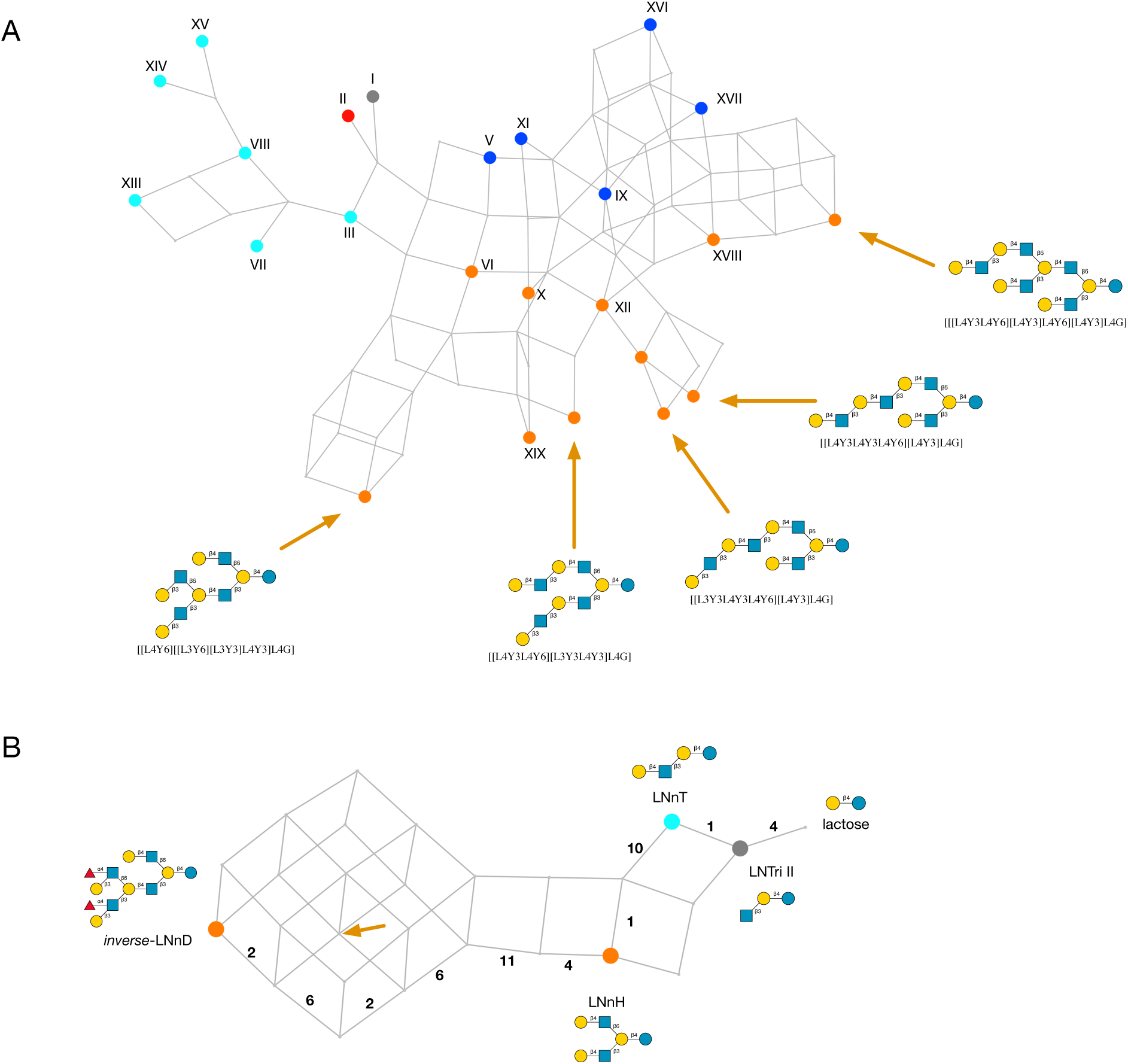
Biosynthesis of HMO core structures. **A**. Network of HMO core structures, labelled I–XIX [23] and with five newly identified cores indicated by arrows, with their Glycologue structural identifiers. **B**. Proposed biosynthetic network from lactose to *inverse*- LNnD [55], labelled with observed intermediates. Multiple routes to the same product are shown, with the enzyme numbers labelling the edges (reactions) of one possible route. The position of the novel HMO core structure is shown by the orange arrow. Nodes are coloured according to the base-core code of Fig. 4.

The only omission from the network is core IV, which was not predicted (see Discussion, Other enzyme activities). Given that model predicts a subpopulation of 1/1 and 1/2 base-core structures, which are not part of the observational dataset, of interest are the existence of the “delayed branching” structures, *novo*-LNnO [23, 56], which is Core XIII [23], and its monofucosylated derivative, F-*novo*-LNnO [23], since they display the 1/1 branching pattern elsewhere in the molecule. An additional core structure, not included in the HMO library, is F-*novo*-LNO, which is type 2 on the upper and type 1 on the lower arm of the branch, and 4- α-fucosylated on the GlcNAc of the terminal lacto-*N*-biose [56]. All three HMOs are predicted by the model, which are defined as 0/2 according to the base-core classification. Their structure identifiers, trivial names and simulated biosynthetic enzyme sequences are:

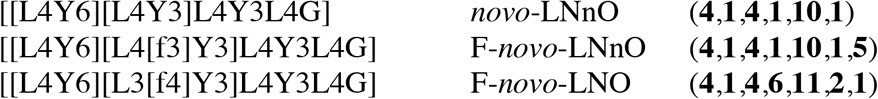

In each case, cIGnT or dIGnT acts on [L3Y3L4Y3L4G] (*para*-Lacto-*N*-neohexaose; Core VII) or [L4Y3L4Y3L4G] (*para*-Lacto-*N*-neohexaose; Core VIII), respectively. The question remains as to the specificity of the 3-β-galactosyltransferase enzyme responsible for extending the 6-linked GlcNAc, whether it is blocked when that residue is immediately connected to lactose, viz., [[Y6][L3Y3]L4G] and [[Y6][L3Y3]L4G].

### Biosynthesis of *inverse*-LNnD

A biosynthetic pathway of the difucosylated *inverse*-LNnD [57], with structure identifier [[L4Y6][[L3[f4]Y6][L3[f4]Y3]L4Y3]L4G], and composition H6N4, is shown in Fig. 5B. It requires the actions of the two GalT enzymes, and is the product of a set of possible sequences of activities, such as (**4**,**1**,**10**,**1**,**4**,**11**,**6**,**2**,**6**,**2**), when acting on lactose as initial substrate. It is based on lacto-*N*-neohexaose, LnNH, a core-2/2 structure that has been reported in several studies [31, 32, 36, 37], and which is itself derived from the core-0/2 lacto-*N*-neotetraose (LNnT) [31–34, 36, 37]. Of note is the (**4**,**11**,**6**) subsequence that is indicative of the formation of the 1/1 motif, even though its base core is 2/2 (cf. Fig. 1: Base core patterns). In Fig. 5B the position of the novel core structure, with Glycologue identifier [[L4Y6][[L3Y6][L3Y3]L4Y3]L4G], is marked by an arrow.

### HMO compositions vs. structures

As a test of the model, we considered the monosaccharide composition of HMOs as a crude grouping of structures. We entered the compositions of the sub-population defined above into GlyConnect Compozitor [58, 59], a tool that roughly simulates the incremental addition of monosaccharides from a nucleotide sugar to an acceptor. The network of compositions is shown in Fig. 6 where we use the condensed notation (hexose=H, hexosamine=N, fucose=F, sialic acid=S and sulfate=s).

**Figure 6.**
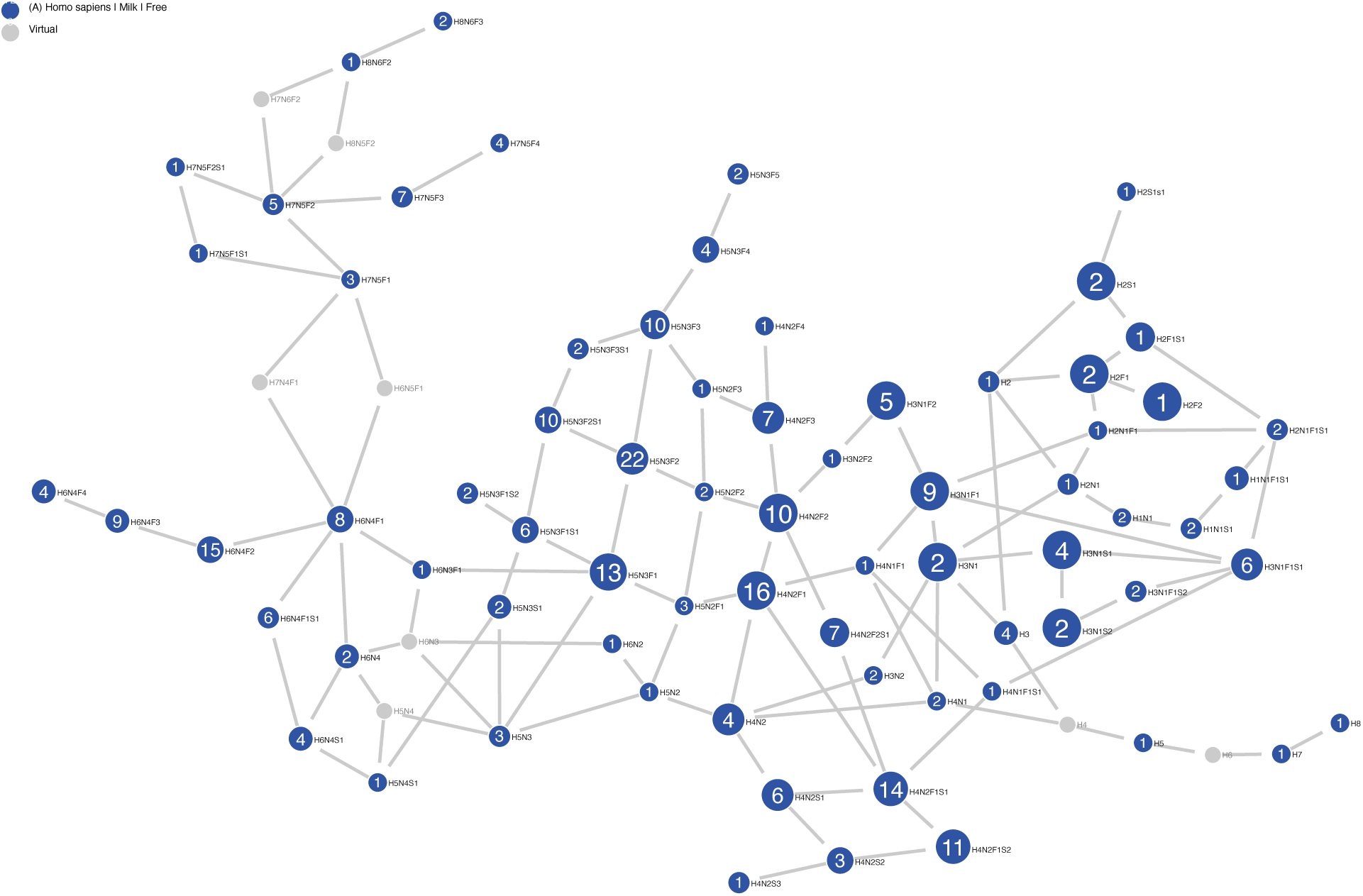
Network of HMO compositions. Nodes represent distinct compositions of hexose, *N*-acetylhexosamine, fucose and sialic acid residues, and differ by one monosaccharide unit from each of their nearest neighbours. Numbers on blue nodes refer to the number of structures in GlyConnect [60] with the composition. Virtual nodes (grey) represent unknown structural intermediates.

In this process, missing intermediates are inferred as virtual nodes depicted in grey. The HMO structures of our library represent 69 compositions stemming from lactose (Hex_2_=H2) (Supplementary Fig. S2) and three from LacNAc (H1N1)) (Supplementary Fig. S3). In these Supplementary figures, the full network is shown with paths highlighted in orange. This colouring is triggered by hovering the mouse on a node to visualise the reachability of and to other nodes from this point (incoming arrows in orange, outgoing arrows in turquoise). The size of the node reflects the number of publications confirming the presence of the corresponding composition. Candidate structures matching a given composition are suggested as part of the library described above that is stored in GlyConnect [60] and accessible in the dedicated HMO section (https://glyconnect.expasy.org/hmo).

We examined the missing compositions as revealed by the virtual grey nodes, especially those with the greatest influence on the connectivity of the graph. In particular, the absence of intermediary structures between H6N4F1 (matching 8 known structures) and H7N5F1 (matching 3 known structures) led Compozitor to consider H7N4F1 or H6N5F1 as potential connectors. Clearly, if not for those virtual nodes the graph would be disrupted. In the same way, missing data connecting H7N5F2 (matching 5 known structures) and H8N6F2 (matching one known structure) are proposed as H7N6F2 or H8N5F2.

The model validated H6N5F1, as 925 generated structures matched this composition, whereas none were associated with H7N4F1. The enzymes of Table 2 can be divided into subsets based on the type of monosaccharide being transferred. Thus, the set of hexosyltransferases are H_E_={**1**,**6**}, *N*-acetylglucosaminyltransferases are N_E_={**4**,**10**,**11**}, fucosyltransferases are F_E_={**2**,**3**,**5**} and sialyltransferases are S_E_={**7**,**8**,**9**}. The graph indicates a preference for GlcNAc first and Hexose second, which could be explained by the mutual interplay of subsets H_E_ and N_E_, each of whose members, in our model, act on the products of the other. Starting from lactose (H2), we expect members of N_E_ to act first, followed by those of H_E_, giving H(2+i)N(i) as the expected composition pattern of cores. By the same reasoning, the we would expect a route from H7N5F2 to H8N6F2 to pass through virtual node H7N6F2 (N+1, H+1), in preference to H8N5F2 (H+1,N+1). This was verified by simulations, which predicted 12955 of the former composition, but none of the latter.

Two virtual nodes represent the intermediary linear structures between existing H3, H5 and H7. These are based on preliminary structure assignments from a set of HMOs not included in our validation set, and which are likely to be novel linear chain polygalactosyllactoses (see Discussion, Other enzyme activities).

The last two virtual nodes are redundant, in the sense that their removal would not disrupt connectivity of the network. However, in their presence, H6N4 (matching 2 known structures) is reachable from 14 nodes in the network and from lactose in particular (blue outgoing arrows in Supplementary Fig. S4). When virtual nodes are not considered then H6N4 is not only reachable through H6N3 or H5N4 and therefore not from the original lactose.

The dataset included in [61] contains 102 HMO compositions some of which representing unexpected extra-large HMOs with a mass greater than 4 KDa. This larger dataset was compared to our library revealing a 70% overlap (Supplementary Fig. S5). Disregarding the extra-large HMOs, 95% of additional compositions were connected to the main network via virtual or existing nodes. Fourteen virtual nodes were required to complete which for most are new in comparison to the six virtual nodes needed in our library.

### Epitopes

HMOs display a wide variety of epitopes, as shown in Fig. 7. The expression of antigenic determinants on HMOs is a function of genetic blood group type and the secretor status of the mother [62–65]. In general, the simulation output matched the percentages of structures with each epitope, except for the Lewis (S)Le^x^ and Le^y^, which are predicted at higher proportions than in the HMO sample library. Since the model lacks an α3GalNAcT enzyme, the only A-antigen bearing HMO, A-hepta [37], was not predicted.

**Figure 7.**
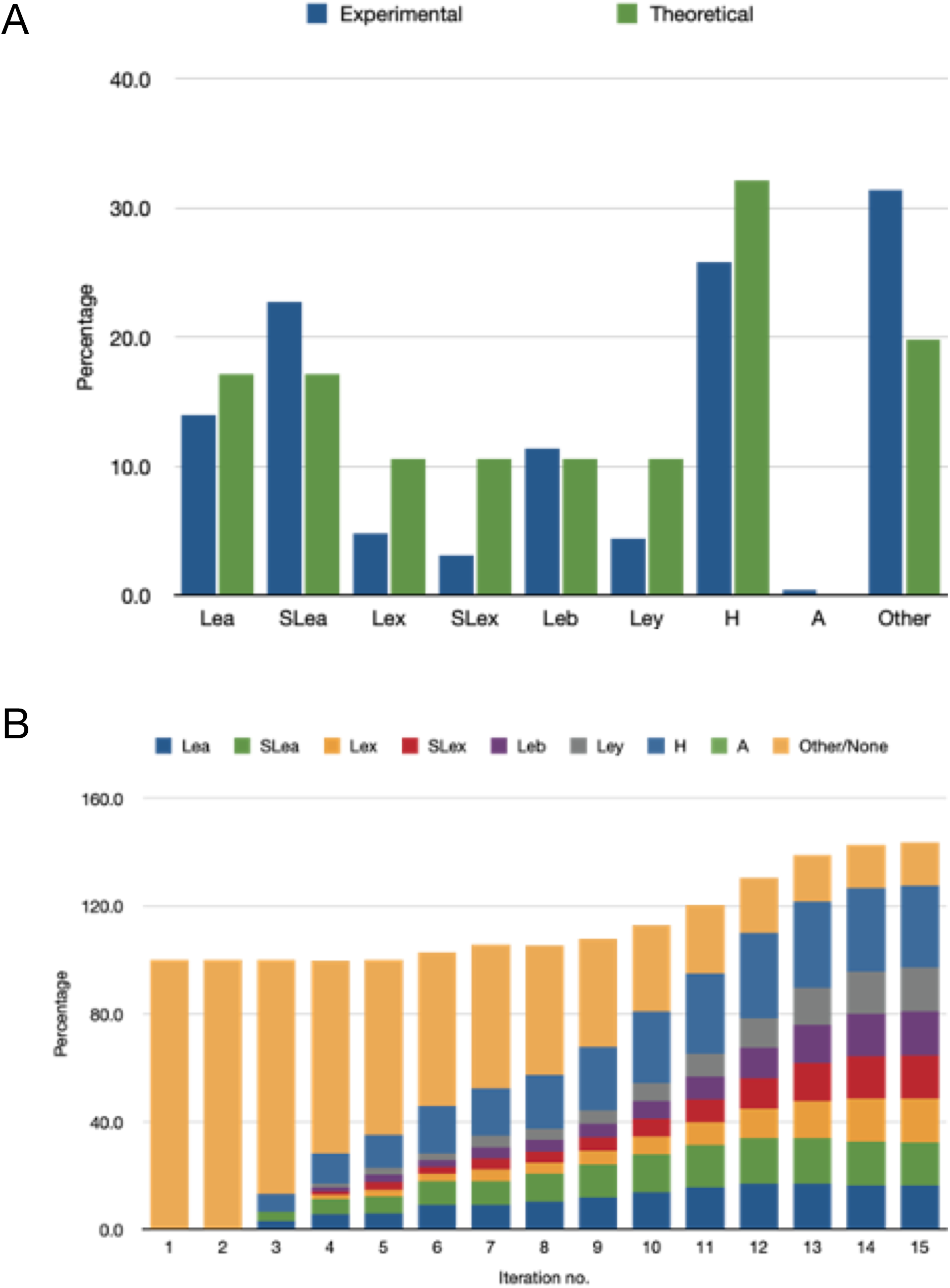
Numbers of epitopes appearing in HMO populations, real and simulated, expressed as a percentage of the total number of structures. **A**. Percentages of the total numbers of each epitope in the experimental (*N*=226) and simulated (*N*=51982; *i*=13) HMO populations. As a result of multi-antennarity (branching), more than one epitope can be present on a single HMO. **B**. Percentages of epitopes appearing in simulations, to which a limit of four GlcNAc residues per HMO was applied, as a function of iteration number (*i*).

Lewis b occurs less frequently on the lower arm of an I-branched type-2 structure, a motif observed in one HMO by Remoroza et al (2018) [32], although not by Wu et al. (2010) [37]. Several examples occur among the HMOs synthesised by Prudden and co-workers [31].

### Enzyme knockouts

The effect of knocking out enzyme activities *in silico*, on coverage of the HMOs in the sample library was investigated. With the null knockout represented by **0**, each activity, **1**–**11** of Table 2 was disabled in conjunction with another, to form single (***x***/**0**) or dual (***x***/***y***) knockouts. The results are summarised in Fig. 8A, in which it is evident that the null knockout, (**0**/**0**), resulted in the highest coverage of the sample data (91.1%), while the other knockouts decreased the coverage to varying degrees.

**Figure 8.**
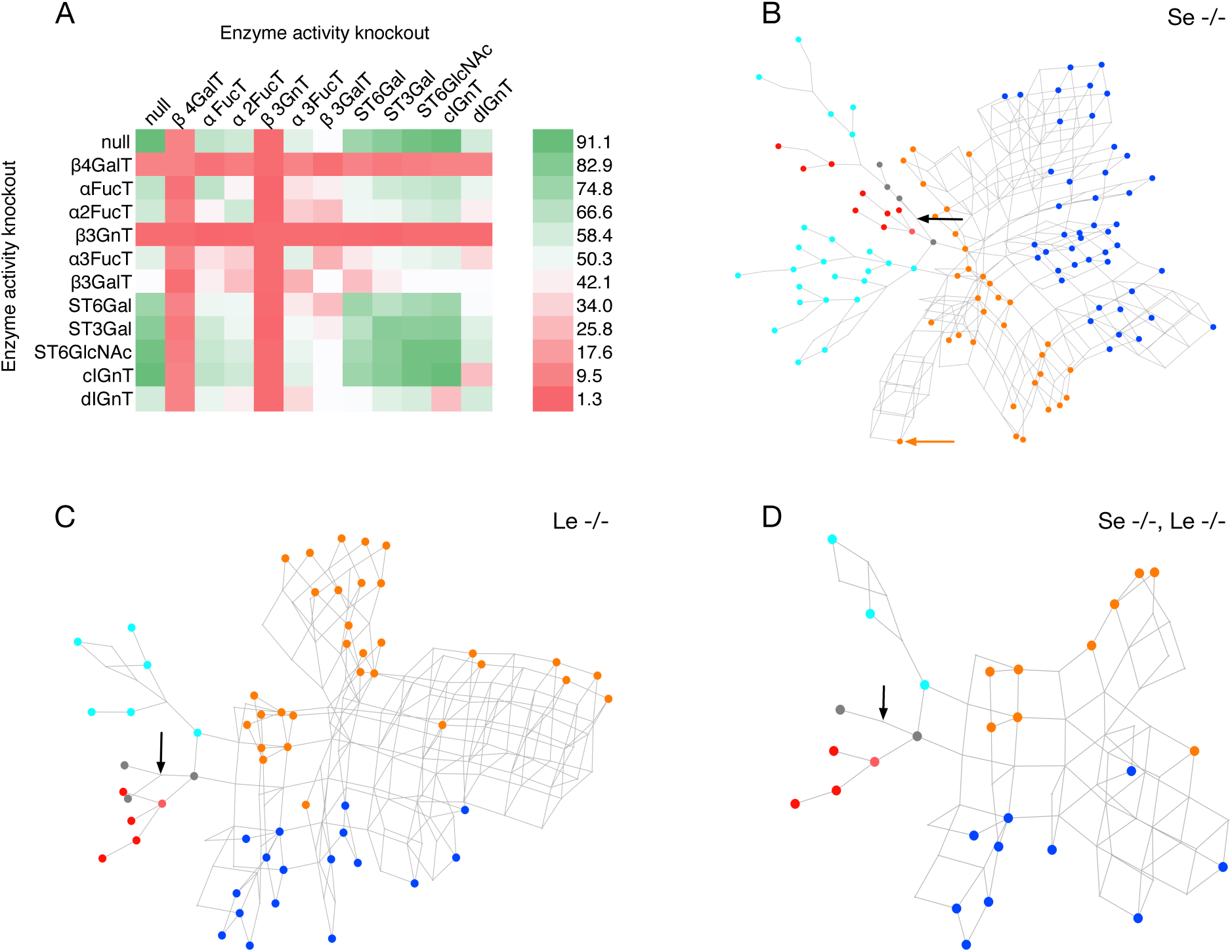
Simulated knockouts of the enzyme activities. **A**. Effects of dual knockouts of the activities in Table 2 on the percentage coverage of the HMO structure library (Supplementary Table S1). In the null/null knockout all enzymes remain active. Colours range from purple (minimal coverage: 1.3%) to yellow (maximal coverage: 91.1%), corresponding to the null/null knockout (**0**/**0**) and the α4FucT/β3GnT knockout (**2**/**4**), respectively. **B**–**D**. Predicted HMO biosynthetic networks of (B) non-secretor (Se-/-), (C) Lewis-negative (Le-/-) and (D) non-secretor/Lewis-negative mothers. Observed HMOs are coloured according to their base-core value (Fig. 1): magenta (1/1), blue (2/1), green (1/2), orange (2/2), red (1), cyan (2), grey (other/unclassified). In each network the position of lactose is indicated by a black arrow. In the Se-/- network (B), an orange arrow indicates the position of *inverse*-LNnD.

Key glycosyltransferases involved in heterogeneity of HMOs are β4GalT (**1**) and β3GnT (**4**), which when knocked out individually resulted in the lowest coverage values consistently.

The lowest coverage, of 1.3%, was obtained with the dual knockout of the α4FucT and β3GnT (iGnT) activities. The distal IGnT activity (**11**), having a broader specificity than the central IGnT (**10**), has the greater influence on heterogeneity. The enzymes ranked according to their influence on overall overage of the HMO sample population (the diagonal elements of the square matrix in Fig. 8), are

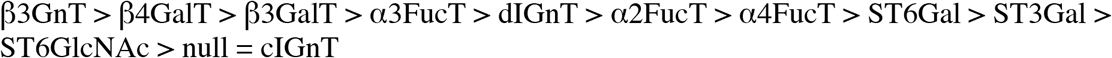

where A > B results in a greater reduction in coverage of the library population when enzyme A is knocked out, compared to the knockout of enzyme B. Maximal coverage was attained with the control (null) knockout, equal to that of the cIGnT knockout.

HMO antigens vary widely between different human populations, but are broadly categorised according to the secretor (Se) and Lewis (Le) status of the individual. Minimal biosynthetic networks corresponding to non-secretor and/or Lewis-negative mothers were constructed by simulating FUT2 and FUT3 genetic knockouts, corresponding to the elimination of 2-α- and (3/4)-α-L-fucosyltransferase activities of Table 2. A single knockout of α2FucT (**3**) removes all H antigen, and Le^b^ and Le^y^ epitopes; the resulting network, with all other enzymes available, is shown in Fig. 8B. A FUT3 knockout was modelled through elimination of both the α3FucT and α3FucT activities (enzymes **2** and **5**), with the result shown in Fig. 8C. A still smaller population of HMOs is predicted when all three fucosyltransferase activities were removed (Fig. 8D).

## Discussion

We have shown that 11 enzyme activities of the model can account for more than 90% of all HMOs analysed in this study, for which we have computed possible reaction networks. We proposed a biosynthetic pathway leading to *inverse*-LnND, which has revealed a novel HMO core, one of five such cores disclosed by this study. Linking composition data to enzyme activities has enabled us to discriminate between possible routes, where unknown intermediates are inferred to exist. While the HMO-Glycologue simulator can be tailored for 2^11^ possible phenotypes, the single- and dual- knockouts of enzyme activities enabled us to rank the enzymes according to their degree of influence on the observed HMO population. The influence of β3GnT and β4GalT activities on heterogeneity is not unexpected, as both are involved in the extension of oligosaccharides via LacNAc repeats.

### Kinetic and genetic regulation of HMO core types

That the majority of branched HMOs fall into the two 2/1 and 2/2 base-core categories could be explained by the higher activity of EC 2.4.1.38 (**1**) towards GlcNAc-β1,6-Gal than to GlcNAc-β1,3-Gal [66], which would suggest that a kinetic competition exists between the two galactosyltransferases (**1**,**6**) that favours type-2 termination on the 6-linked arm. This would also help to explain the asymmetry in the existence of LacNAc repeats among the sample population, which appear only on the 6-linked GlcNAc. Of the 226 HMOs in Supplementary Table S1, there are no occurrences of LacNAc-extended (type-2) terminations of “lower” branch and three occurrences of lacto-*N*-biose extended type 1; of the “upper” branch, there are 28 of lacto-*N*-biose extended (type 1) and 53 of LacNAc extended (type 2). Since changes to HMO concentration are observed to vary over the course of lactation [33, 67, 68], owing to regulation of the expression of the genes coding for these enzymes, as well as their location in the Golgi, a spatiotemporal separation of the different galactosyltransferase activities **1** and **6** may account for the synthesis of *inverse*-LNnD (Fig. 5).

### Non-predicted structures

The 20 HMOs of Supplementary Table S1 that were not predicted by the model form a set of structures that may point, either to novel enzyme activities unique to milk, or else to alternative functions of known enzymes. In the following, we discuss possible routes of formation of these unknowns in terms of known, and possibly novel, enzymes. The structures and reactions are summarised in Fig. 9.

**Figure 9.**
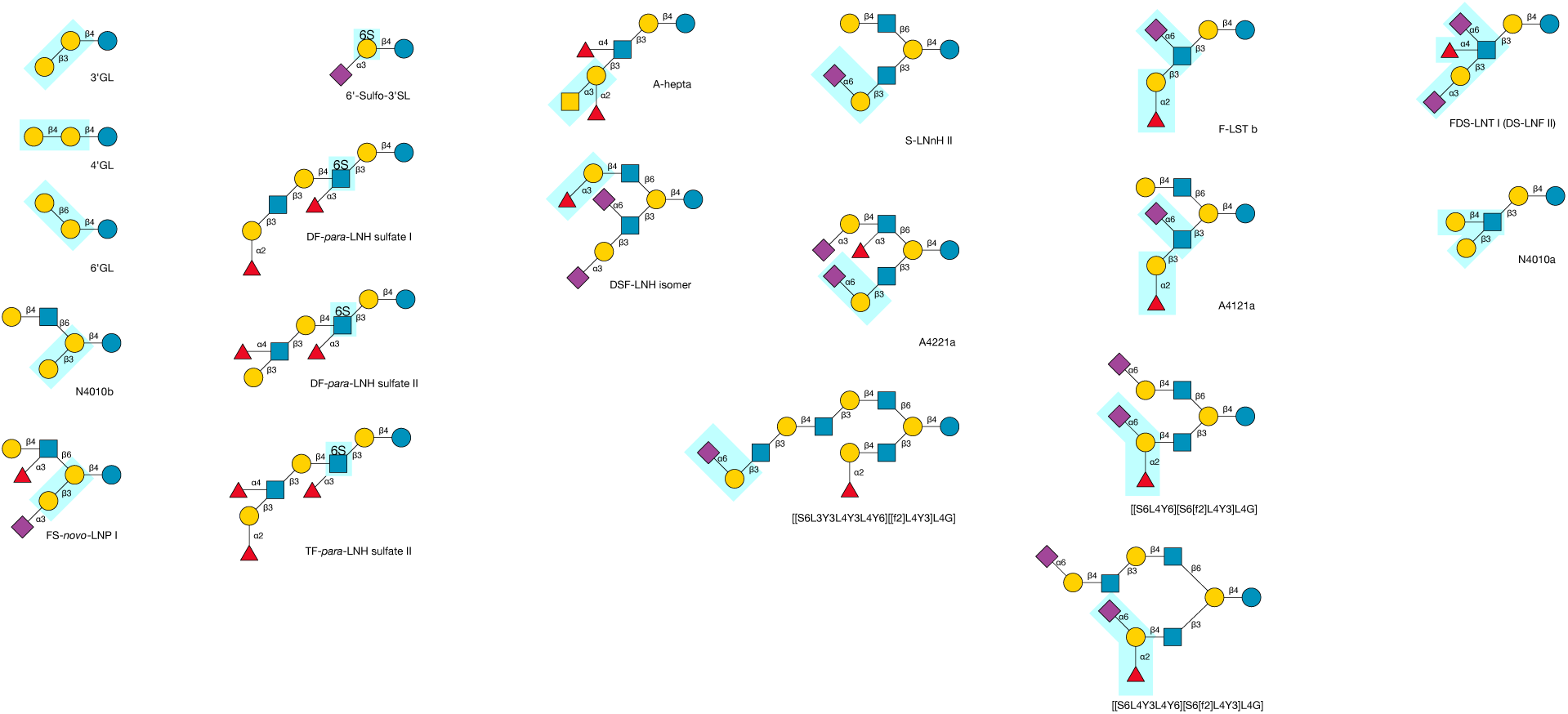
Structures of HMOs that were not predicted by the model. Highlighted motifs (boxed regions) suggest possible additional enzyme activities. See text for details.

### Galactoside β-galactosyltransferases

Three galactosyllactose structures [L3L4G] (3’GL), [L4L4G] (4’GL) and [L6L4G] (6’GL) were found. The presence of 6′-galactosyllactose in the protein fraction of human milk from non-secretor mothers was first reported by Yamashita and Kobata [69], who also demonstrated that it was the product of a specific galactosyltransferase and not a transgalactosidation reaction of β-galactosidase. Two further galactoside β- galactosyltransferases acting on the 3’ and 4’ positions of lactose are necessary to explain the other two non-predicted structures in this subset. No other extensions of the galactosyllactoses were found in this study, though linear chain poly-galactosylated HMOs occur in the milks from other species, such as lion [70].

### GlcNAc β-1,4-galactosyltransferase

The structure of [L3[L4]Y3L4G] (N4010a [32]) is unusual in possessing both type 1 and type 2 termination. While it could be represented within our model in three different ways (see Methods), including as [L4[L3]Y3L4G], no LacNAc extension of LNnT was observed with 3-β-galactosylation of the GlcNAc, therefore we propose that the parent is lacto-*N*-tetraose ([L3Y3L4G], LNT), decorated by an unknown GlcNAc β-1,4-galactosyltransferase. The Glycologue structure identifier thus acts as a record of distinct enzyme activities, even where the structures are otherwise indistinguishable.

### Alternative sialyltransferase activities

Based on the activities of other sialyltransferases, which act after the fucosyltransferases to cap Lewis-type termini, the ST3Gal and ST6GlcNAc activities of the model could be modified similarly to account for the non-predicted structures, [[f2]L3[S6]Y3L4G] (F-LST b) and [S3L3[S6][f4]Y3L4G] (DS-LNF II). By a similar modification the 6-sialylated H2- antigenic structures, [[S6L4Y6][S6[f2]L4Y3]L4G] and [[S6L4Y3L4Y6][S6[f2]L4Y3]L4G], of Prudden et al. [31] could be modelled.

Although the 6-sialyltransferase activity of EC 2.4.99.1 (**7**) is generic, acting on β-galactosyl termini, we modelled its major activity, which is towards the type-2 LacNAc acceptor [64]. The enzyme from goat and bovine colostrum has a secondary activity towards type 1, which might explain the three HMOs in which this motif appears, [[L4Y6][S6L3Y3]L4G] (S-LNnH II) [32, 38], [[S3L4[f3]Y6][S6L3Y3]L4G] (A4221a) [32] and [[S6L3Y3L4Y3L4Y6][[f2]L4Y3]L4G] [31].

The disialomonofucosyllacto-*N*-hexaose structure, [[[f3]L4Y6][S3L3[S6]Y3]L4G], listed by Smith et al. [34] could not be predicted owing to the lack of a 3-α-fucosyltransferase acting on galactose. The corresponding 2-α-fucosylated structure, variously known as FDS-LNH I [30] and DSF-LNH II [23, 32] is predicted within the system described here.

### 6-*O*-sulfotransferases

Only 4 of the 226 HMOs in the library were sulfated, with structure identifiers [S3[s6]L4G], [[f2]L3Y3L4[f3][s6]Y3L4G], [L3[f4]Y3L4[f3][s6]Y3L4G] and [[f2]L3[f4]Y3L4[f3][s6]Y3L4G]. We conclude that there exist two, as-yet uncharacterised, 6-*O*-sulfotransferase enzyme activities of human milk, which are specific for lactose and LNTri II, since no other sulfated compounds were found. All four non-predicted sulfated HMOs have non-sulfated counterparts that were predicted by our model.

### Other enzyme activities

The existence of A-hepta, discovered by Wu et al (2010) [37], is likely to be the product of EC 2.4.1.40, glycoprotein-fucosylgalactoside α-*N*-acetylgalactosaminyltransferase, acting on the H antigen. Since only one occurrence of the antigen was found in the sample dataset, this enzyme activity was excluded from the model.

Following the publication of a mass spectral reference library for HMOs based on the NIST Standard Reference Material (SRM) 1953 dataset [32], the structures of several unknown HMOs were inferred from the original library by means of a bootstrapping approach [70]. Out of 78 of the HMOs newly identified (Table S4 of [70]), 48 (61.5%) were predicted by HMO-Glycologue with the 11 enzymes of the current model active. Since many of the structural assignments are preliminary, and coverage significantly lower than for each of the individual we did not include these data in our library (Supplementary Table S1). Nevertheless, a review of those structures which were not predicted is instructive, since the existence of some of them might be explained by our current knowledge of the enzymes of human glycosylation. In what follows, R*n* refers to HMO with index number *n* within the cited dataset [70].

HMOs R1 and R5, Galactosyl-FpLNnH (Glycologue structure identifier: [[L4Y3L4[f3]Y6][L3]L4G]) and Galactosyl-TFpLNH ([[[f2]L3[f4]Y3L4[f3]Y6][L3]L4G]), could be formed from or more of the *O*-glycosyltransferase core enzymes, if these were active against lactose, to give [[L3]L4G] followed by [[Y6][L3]L4G]. The corresponding GalNAc-linked glycan, [L4Y3L4[f3]Y6][L3]VT, is predicted within O-Glycologue. If the two enzymes, C1GalT and C2GnT, were co-expressed in human mammary gland, then such 3-galactosylated “Core-2 HMOs” would be expected to be abundant, yet they are not. The [L3]L4 motif is observed in only one other structure, N4010b, from this and a previous [32] study, although structure R2, Galactosyl-FpLNnO, might be another candidate. Instead, a possible explanation for N4010b, [[L4Y6][L3]L4G], is that it is the product of C2GnT acting on 3′-GL, formed by the galactoside 3′-β-galactosyltransferase referred to above, followed by β4GalT:

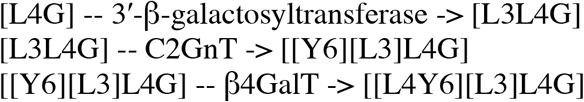

The structure is also known as HMO core IV, lacto-N-novopentaose I [23]. By similar reasoning, Novo-LNP I (R7) could be a “Core-4” HMO. The enzymic origin of these structures therefore remains to be elucidated.

The α-galactosylated structures reported by that study may be the products of two enzymes, fucosylgalactoside 3-α-galactosyltransferase (EC 2.4.1.37) and *N*-acetyllactosaminide 3-α- galactosyltransferase (EC 2.4.1.87). O-Glycologue [71] currently models the activity of the former, EC 2.4.1.37, and thus could be modified to predict these structures. The activity of EC 2.4.1.87 could likewise be added; judging by their absence from this dataset, this enzyme seems not to be active toward fucosylated HMOs.

In this context, our analysis of HMO compositions, our results concluded that certain virtual nodes were predicted within the system, but not others (see HMO compositions vs. structures). Interestingly, composition H8N5F2 that is virtual in our library network is real in the Porfirio et al. [61] dataset, thereby supporting the selection of H8N5F2 over H7N6F2 as a valid path connecting H7N5F2 and H8N6F2. If α-galactosyltransferase enzymes are active during lactation, it might also explain the higher ratio of hexose to GlcNAc of such compositions.

The dodecaose series [70] are differentially fucosylated, and all of them would be predicted by the model if it were not for the presence of the repeating L3Y3 units (L3Y3L3Y3) on the lower branches of around half of them. This unusual sequence, reported in *O*-glycans of human colonic [72, 73] and gastric [74] mucin, are not formed by EC 2.4.1.149, which is instead responsible for the formation of polyLacNAc. Their presence would suggest the activity of an unknown β3GnT enzyme activity that recognises only the terminal [L3Y3, as acceptor, since the type-1 extended upper arm, L3Y3L3Y6, is not observed. Another sequence that is novel is L4Y3L3Y3, i.e. type-1 structures extended by LacNAc to form type 2, which would infer that β4GalT does not recognise the [Y3L3. We note that the same motif exists in DF-para-LNO I (R58), to which the structure identifier [[f2]L3[f4]Y3L3Y3L4Y3L4G] was assigned.

The novel Iso-LNnO oligosaccharide would require a β6GalT enzyme that would form [L6]Y. The β(1->6)-galactosyltransferase referred to above, which forms 6′-GL, might also be responsible for this structure, although, to our knowledge, it remains to be determined if is also active against LacNAc.

The polyGal structures pentagalactosyllactose, heptagalactosyllactose and octagalactosyllactose R71–R73; might be initiated by the enzyme responsible for early heparan/chondroitin biosynthesis, galactosylxylosylprotein 3-β-galactosyltransferase (EC 2.4.1.134), if that enzyme were active towards glucose in addition to xylose. If the di-, tri- and tetra-galactosyllactose precursors of these larger structures exist, they were not reported, nor included in the NIST mass spectral HMO reference library.

## Conclusions

As samples produced in various geographical, nutritional and health conditions accumulate, it is very likely that more HMOs will be identified. In light of the results presented here, we envisage that the Glycologue HMO-enzyme simulator will be extended, or adapted, as our knowledge of the enzymes and their substrate specificities improves. Our attempt to account for the largest collection of currently known HMOs in our model has raised several questions on individual and combined enzyme activities, some of which remain open. It is possible that the unknown I-branching enzyme, dIGnT, is less specific than cIGnT, and that it can use either type-1 (β3-Gal) or type-2 (β4-Gal) HMOs as acceptors. The removal of cIGnT, should its activity turn out to be redundant is an option. It is hoped that this analysis will promote further examination of these enzymes, including those yet to be characterised, such as the galactoside β-galactosyltransferases. With its predictive power, the model can also be considered as a guide for experimental synthesis of HMOs, which would potentially enable testing with specialised microarrays.

## Methods

The method is based on a formal language, which uses a single-letter code for the monosaccharides, as shown in Table 1, and a set of transformation rules that add one monosaccharide at a time, using a regular-expression based pattern matching to model the enzyme activities. A software implementation of the method, Glycologue, acts iteratively on an initial acceptor-substrate, passing it to each enzyme in turn, and accumulating a set of acceptor-products. The pool of novel acceptor-products become the substrates at the next iteration, until either no new products are formed, or a maximum number of iterations set by the user has been attained. Since extension of oligosaccharides occurs principally by means of LacNAc (Gal-β1,4-GlcNAc), simulations could be limited by the number of GlcNAc residues incorporated. The reaction network was deemed to have *closed* at iteration *i* when no further products were added at the next iteration, *i*+1. Simulations could also be limited by supplying a target composition value such as H4N3F1S1 (4 Hex, 3 HexNAc, 1 x dHex, 1 x Neu5Ac), such that enzymes would act on a substrate if and only if its composition did not exceed the prescribed value of any class of monosaccharide. The number of possible oligosaccharide structures matching that composition was then calculated.

Table 2 lists the biosynthetic enzymes predicted by the model, with an index number, **1**–**11**, the EC number, where available, a short name, a longer accepted name and a reaction pattern. The reactions in Table 2 are based on activities of enzymes already classified within the IUBMB Enzyme List, or from the cited references, wherever an EC number is not available. In reaction patterns, asterisks act as a wildcard character, to denote portions of the molecule of indeterminate length. The glossary in Table 2, footnote b, shows some of the assumptions implicit to the model, such as the anomeric configuration of the donors, from which it can be inferred which enzymes invert or retain the stereochemical configuration of the donor during incorporation into the acceptor. Reactions could, in addition, be limited by Boolean conditional regular expressions. In the case of two I-branching enzymes, **10** and **11**, the assumption was made that these enzymes would not be active towards 3-fucolactose (3-FL) substrates.

The types of reaction catalysed are classified according to a limited number of transformation patterns. We consider the default mode of action to be the *extension* of a linear oligosaccharide, by

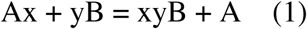

where x and y are monosaccharides, Ax is the nucleotide-sugar donor, and yB the acceptor-substrate.

The formation of a single branch along a linear chain is described as *decoration*, where the pattern is

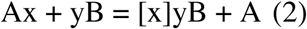

and we have assumed that [x]y is a shorthand for *[x]y*B, the asterisks acting as a wildcard character. Double branches are used to form symmetric core structures, such as the trimannosyl core of N-glycans, or O-linked glycan cores based on GalNAc:

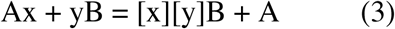

Capping of branches and linearly extended chains is achieved through *termination*, of which sialylation is a typical example. *Modification* of monosaccharides, such as by the actions of sulfotransferases, follows the same pattern as decoration (2). Glycologue structure identifiers order branches by linkage position, writing the branch with the lowest linkage first, reading from right to left. Modifiers are written before sugars units, and multiple modifiers on the same monosaccharide are again ordered by linkage position, from lowest to highest, reading right to left. Boolean conditionals can be applied to enzymes, to prevent action in the presence or absence of a particular recognition motif.

This classification, and the numbering rules, enables the assignment of a unique Glycologue structure identifiers to HMOs with determined structures. Glycosyltransferases **1, 4** and **6** are involved in extension, and the sialyltransferases in termination, by pattern (1); the fucosyltransferases **2, 3** and **5**, and ST6GlcNAc (**9**) are involved in decoration according to reaction pattern (2), and GTs **10** and **11** form double branches according to pattern (3).

The activities of the enzymes could be reversed, and all reaction paths leading from a given HMO to lactose determined. For any set of such initial substrates supplied to the reversed simulator, the complete collection of such paths leading from lactose provided a minimal biosynthetic reaction network. Individual paths are represented by ordered sequences of enzyme activities. For example, the formation of LST b, which has the structure identifier [L3[S6]Y3L4G], has sequence (**4**,**6**,**9**), when starting from lactose. As there can be multiple routes to a given HMO, the networks generated in both the forward and reverse directions possess a lattice-like structure.

A web application interface to the HMO-enzyme simulator, the set of experimentally determined HMOs used as validation and the source code of the simulator as a Python 3 script, are available at https://glycologue.org/m. The Glycologue family of simulators [29, 71, 75] supports import and export of IUPAC short form and GlycoCT condensed formats, along with the native Glycologue structure identifiers, while export as Linear Code is also provided for individual structures. Networks can be exported as SBML, with GlycoCT-XML embedded as annotations of the nodes.

## Acknowledgements

Partly supported by the Swiss Federal Government through the State Secretariat for Education, Research and Innovation (SERI). The Expasy portal is maintained by the web team of the Swiss Institute of Bioinformatics and hosted at the Vital-IT Competency Center This work was part supported by SNSF grant #31003A_179249 (Frederique Lisacek).

## Author Contributions

Writing: AM, FL

Editing: AM, GD, FL

Experiment design: AM, FL, GD

Software development: AM, JM

