## Supplementary Figures for "In silico analysis of the human milk oligosaccharide glycome reveals key enzymes of their biosynthesis"

#### Supplementary Tables

**Supplementary Table S1. Library of 226 experimentally characterised human milk oligosaccharides (HMOs).** Structures are provided as GlycoCT-condensed format and as Glycologue structure identifiers. Core numbers (I–XIX) are those of Urashima et al. (2018) [23]. Compositions are assigned according to the numbers of hexose (H), GlcNAc (N), fucose (F) and Neu5Ac (S) residues per HMO. An online version of this table is available at https://glycologue.org/m/library.php.

**Supplementary Table S2. Human milk oligosaccharide (HMO) core structures.** Structures are provided in GlycoCT-condensed format and as Glycologue structure identifiers. Core numbers listed are those of Urashima et al. (2018) [23; novel cores are as indicated. Compositions are assigned according to the numbers of hexose (H), GlcNAc (N), fucose (F) and Neu5Ac (S) residues per HMO. [Online version at: https://glycologue.org/m/library.php?f=hmo-cores-v1]

### Supplementary Figures

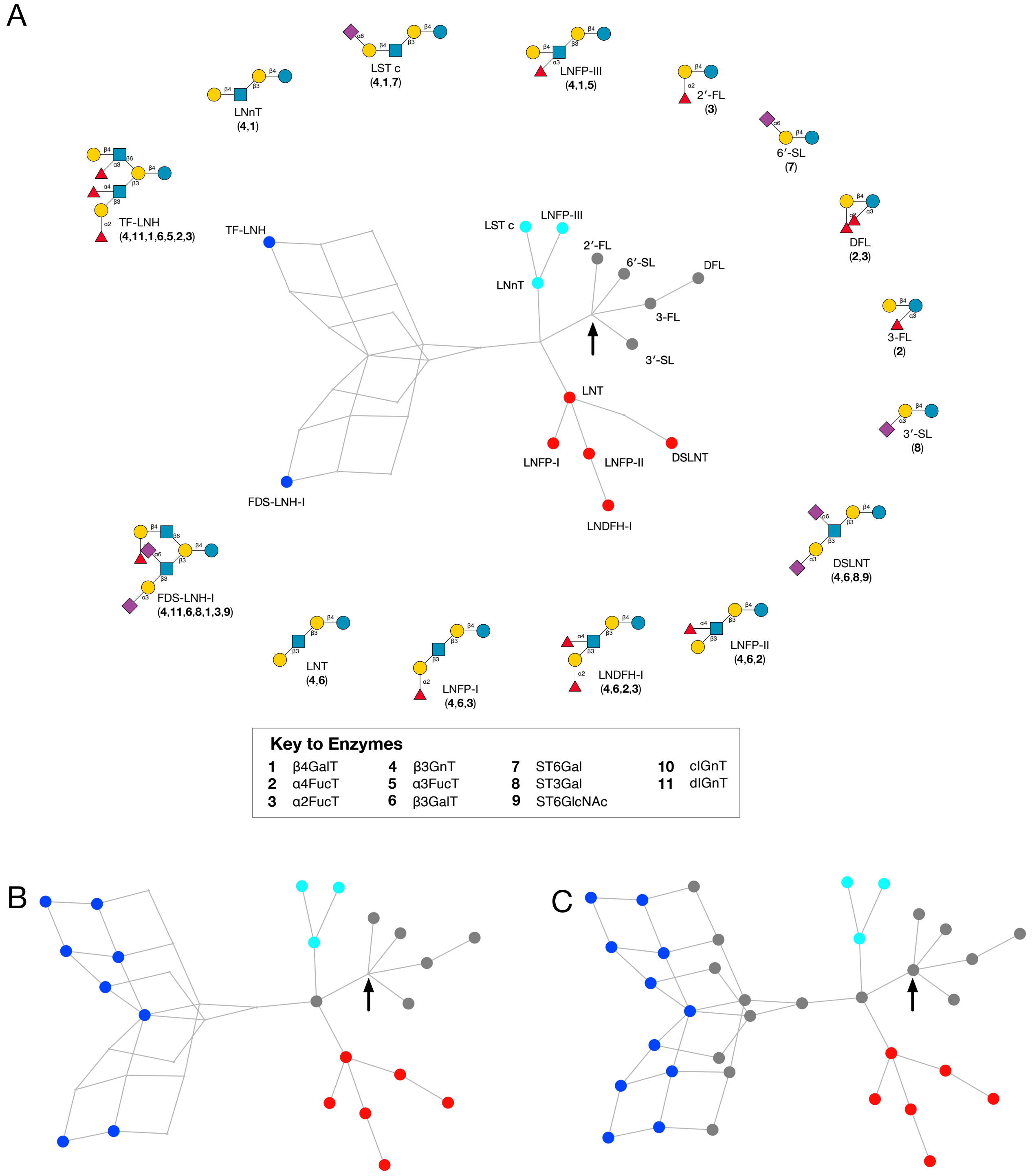


**Supplementary Figure S1.** Proposed biosynthetic network for the 15 most abundant HMOs in mature human milk [54]. Nodes are coloured according to the base core pattern of Fig. 1: cores 2/1 (blue), 0/1 (red), 0/2 (cyan), or other/unclassified (grey). The position of lactose in the network is indicated by an arrow. **A**. The 15 most abundant HMOs are highlighted. Ordered lists of the enzyme activities involved are printed beneath each structure, using the indices shown in the Key to Enzymes (*cf*. Table 2). **B**. As A, with all observed HMOs within the library (Supplementary Table S1) highlighted. **C**. All nodes highlighted and coloured by base core.

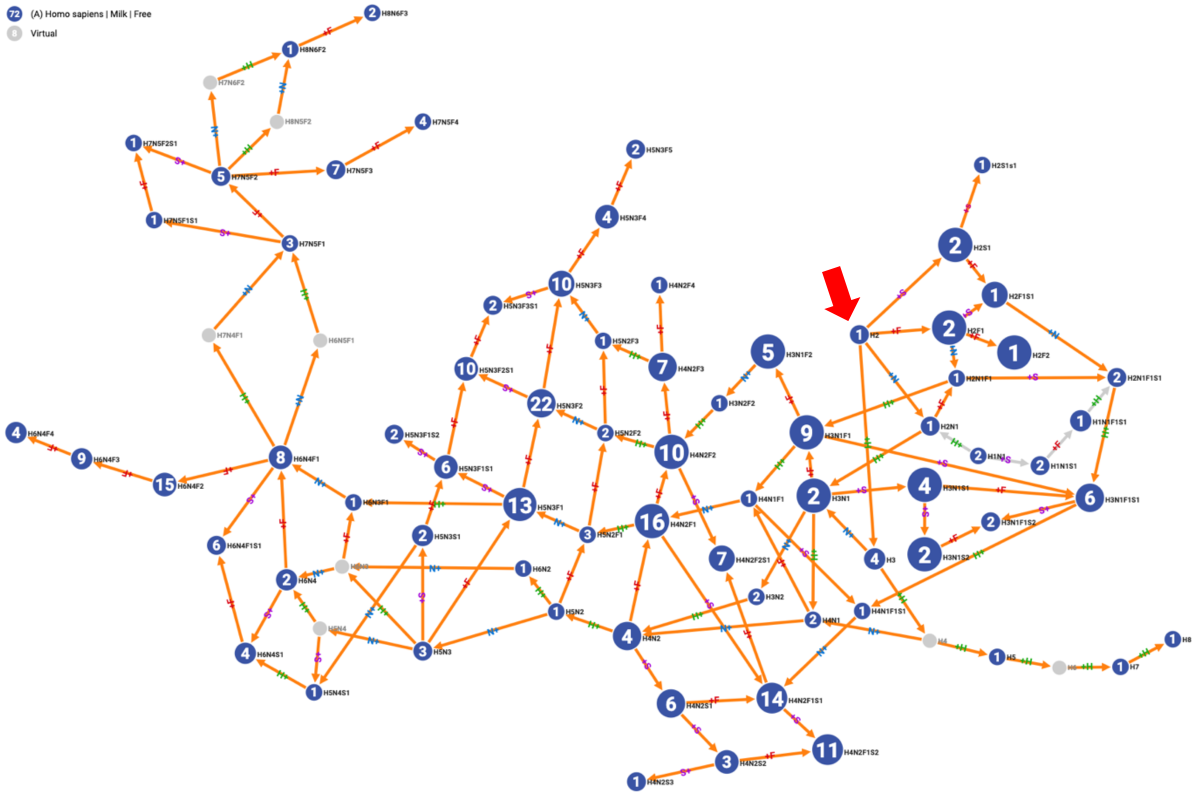


**Supplementary Figure S2.** Graph of HMO compositions highlighting all paths from lactose (red arrow) which shows almost full coverage but not unique LacNAc containing structures.

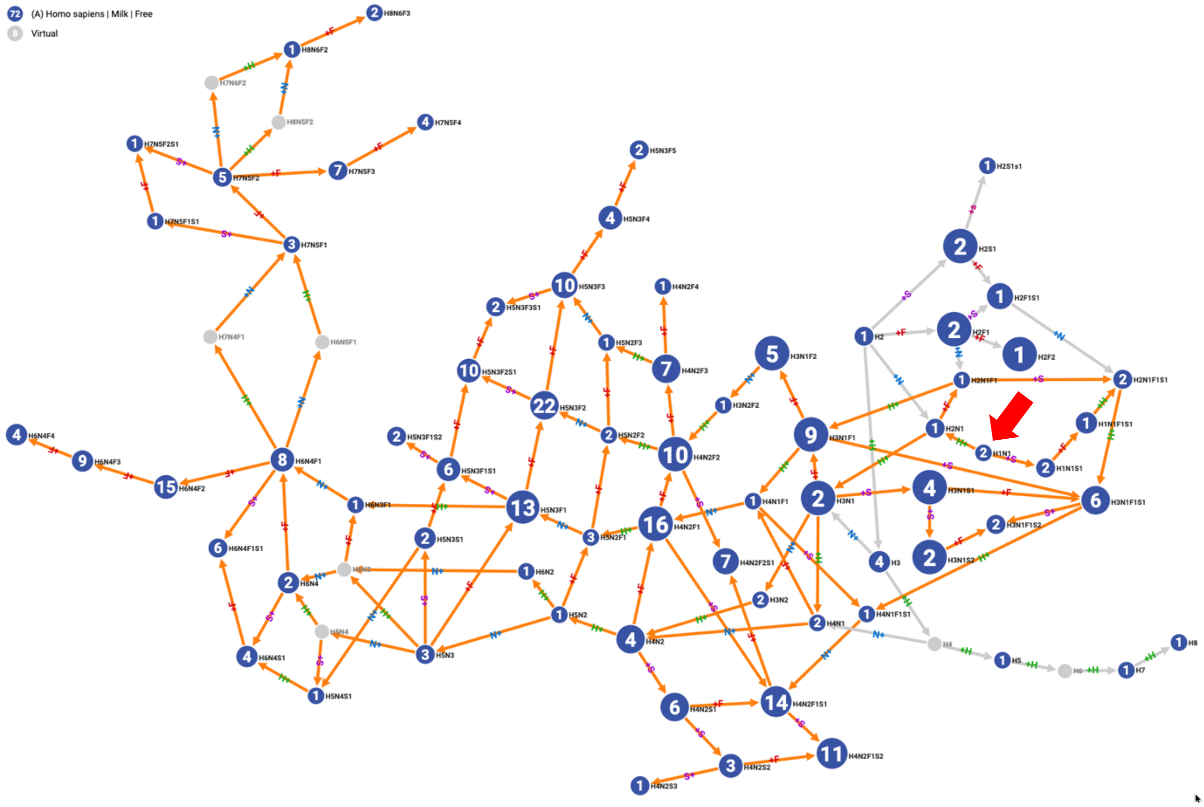


**Supplementary Figure S3.** Graph of HMO compositions highlighting all paths from LacNAc (red arrow) with a lesser coverage – only GlcNAc-free structures.

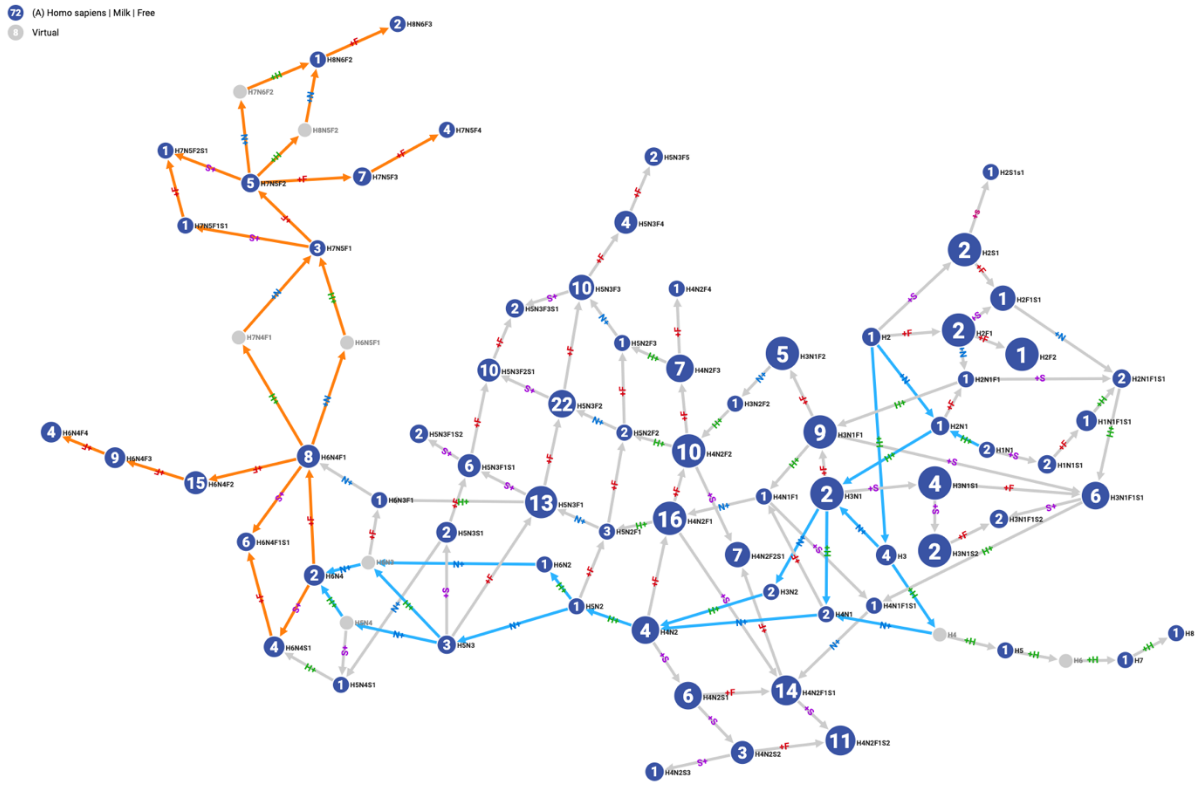


**Supplementary Figure S4.** Graph of HMO compositions highlighting the position of node H6N4 showing that it is only reachable from lactose if virtual nodes (H6N3 or H5N4) are included. If not H6N4 is a terminal node only connecting H6N4F1 and H6N4S1.

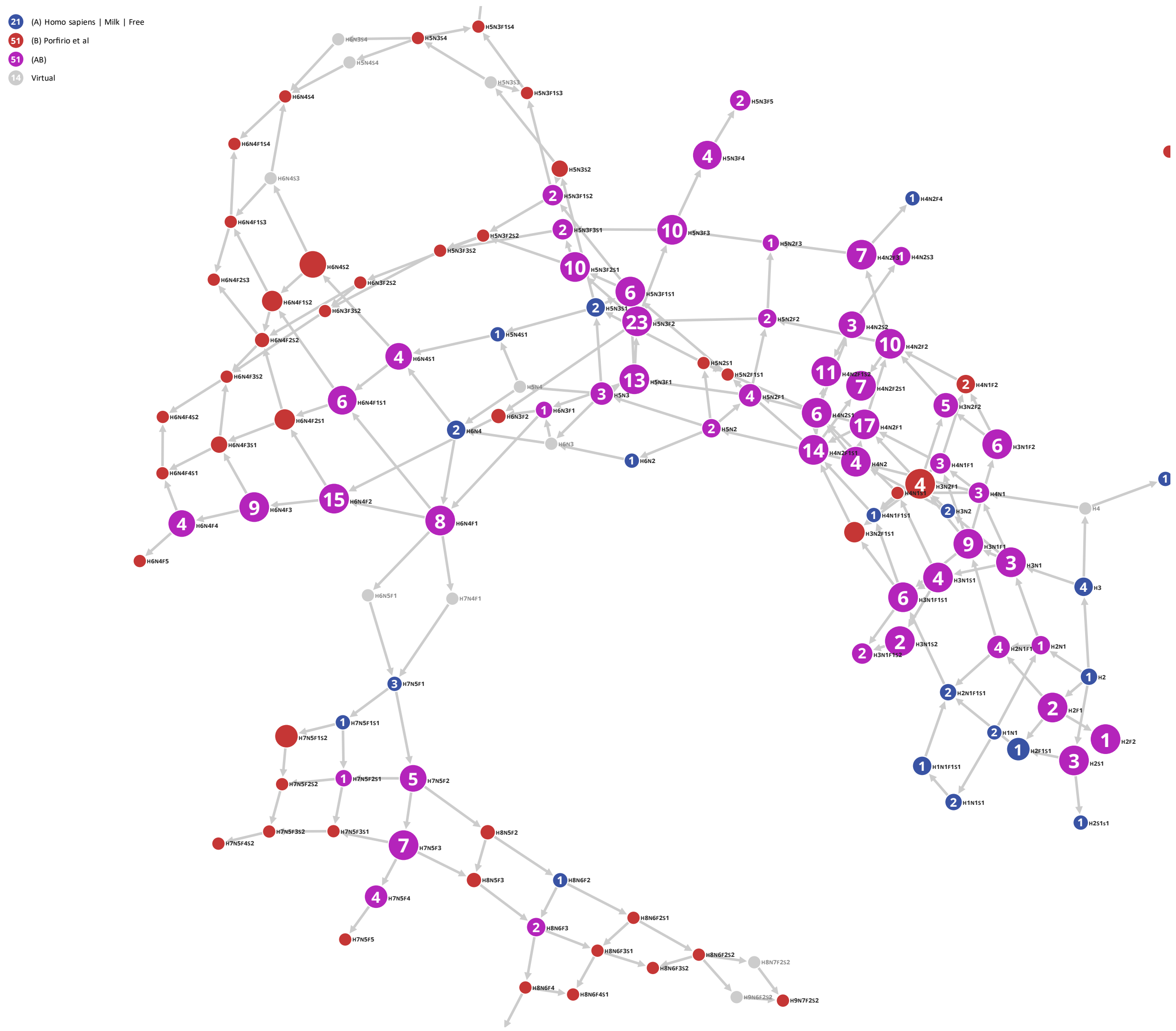


**Supplementary Figure S5.** Composite graph of HMO compositions superimposing the original 72 HMO compositions and the Compozitor output of Porfirio et al. [60] 102 HMO compositions. Magenta nodes are common to both (51/72=70%), blue nodes are specific to the original HMO compositions and red to the (Porfirio et al) data.
